# EnSEMBLE: a framework for enhancer-anchored pathway analysis that locks in enhancer-RNA–corroborated pathways from transcriptome sequencing data for biological validation

**DOI:** 10.64898/2026.08.17.745283

**Authors:** Laizhi Zhang, Akshat Gupta, Yue Wang, Renu Sharma, Bashir Lawal, Gavin Hou, Xiao-Song Wang

**Affiliations:** Department of Human Genetics, School of Public Health, University of Pittsburgh, Pittsburgh, PA, USA; UPMC Hillman Cancer Center, University of Pittsburgh, Pittsburgh, PA, USA; Department of Pathology, University of Pittsburgh, Pittsburgh, PA, USA; Department of Biomedical Engineering, University of Michigan, Ann Arbor, MI, USA

**Author notes:** These authors contributed equally (Laizhi Zhang, Akshat Gupta). Correspondence: Xiao-Song Wang.

## Abstract

**Background:** Pathway discovery methods for transcriptome sequencing return tens to hundreds of redundant gene sets, and biologists often subjectively select the pathways fitting biological expectations. What is missing is not another statistical method, but a way to corroborate each candidate pathway against an independent, mechanistic line of evidence.

**Results:** We introduce EnSEMBLE (Enhancer-Set Enrichment & Mechanism-Based Linked Evidence), a tool that corroborates gene-level pathway enrichment with an orthogonal enhancer layer drawn from the same transcriptome sequencing data: active enhancers transcribe enhancer RNAs already present in standard RNA-seq, so a pathway’s regulatory state can be scored from the very run that produced the gene-level signal, at no added cost. EnSEMBLE pairs pathway enrichments with Enhancer-Program Enrichment Analysis (EPEA), collapses redundant gene sets into process-level Themes, and retains only those that a concordant enhancer program corroborates. This dual-evidence requirement reduced reported signatures by >97% (hundreds of gene sets to 3–18 claims) across four datasets spanning cancer perturbations and iPSC-to-neuron differentiation. Surviving claims recovered expected biology—mesenchymal-program collapse upon *SNAI1* knockout, regulatory convergence during neuronal differentiation—and named mechanisms pathway enrichments missed, including an mTOR–MYC–SPT5 elongation axis in rapamycin-treated PANC1 cells. A language AI agent performs narrative synthesis over deterministic statistics, with reproducibility enforced by temperature-zero inference and three-run consensus. We further provide enhancer over-representation analysis (eORA), mapping non-coding GWAS variants to the same programs to recover cell-type–selective trait associations.

**Conclusions:** EnSEMBLE shifts transcriptomic interpretation from enumerating possibilities to adjudicating evidence, yielding a compact, traceable set of enhancer-corroborated claims that identify the regulatory programs driving cellular change and prioritize them for experimental validation.

## Introduction

### Pathway enrichment has become the bottleneck in transcriptomic discovery

Transcriptome profiling is now a routine readout of cellular state, but interpretation has become the rate-limiting step: a single experiment can yield a large number of differentially expressed genes and pathways, and the field produces such lists far faster than it can explain them. The dominant way to make sense of them is gene-set enrichment analysis, such as Gene Set Enrichment Analysis (GSEA) [1,2], Enrichr [3], DAVID [4], clusterProfiler [5], IndepthPathway [6], etc. Indispensable as they are, these methods are widely recognized to return sprawling, redundant lists of pathways. More troubling, the underlying statistics are fragile: inter-gene correlation inflates significance and false-discovery control is unstable, so the same dataset can yield different “significant” pathways across tools, software versions, and parameter settings [7,8]. Faced with such lists, investigators routinely keep the few terms that fit prior expectation and set the rest aside—a procedure that is subjective and hard to reproduce. What is missing is not another enrichment statistic, but a way to corroborate each candidate pathway against an independent, mechanistic line of evidence.

### Acting on an unverified pathway can cost a year of failed bench work

This gap between statistical enrichment and biological truth is not academic; it sets the pace of discovery. Acting on a pathway that is merely plausible commits months of cell-line work, animal models, and reagents to a hypothesis that may never have been real, and across preclinical cancer biology, a large fraction of such leads fail to replicate when independently retested [9,10]. Our own study of the *BCL2L14-ETV6* fusion, a rearrangement we found to be highly specific to triple-negative breast cancer, is a hallmark case in point[11]. Conventional gene-expression pathway analysis of fusion-expressing cells ranked epithelial-mesenchymal transition (EMT) among the top pathways. Yet nearly a year of work yielded only modest results, until we recognized that the culture system itself lacked the cue needed to activate the EMT program. Once TGF-β and EGF were supplied, the mesenchymal program became unmistakably activated in the fusion-expressing cells. The lesson generalizes: a pathway call worth a year of bench validation must clear a far higher bar than merely ranking among dozens of enrichments—closer to the confidence an experimentalist would demand before committing resources—and gene-expression enrichment alone rarely reaches it. Clearing that bar requires a second, orthogonal line of evidence that reports directly on regulatory state.

### Enhancer RNAs offer an orthogonal, real-time readout of regulatory activity

Finding the true drivers of gene expression requires looking at the “control switches” of the cell: enhancers. Enhancers act as logic gates, combining inputs from transcription factors (TFs) to control genes [12]. Crucially, active enhancers produce non-coding enhancer RNAs (eRNAs), which directly show regulatory activity [13]. Unlike mRNA levels, which take hours to build up and break down, eRNAs appear rapidly before the target gene is turned on, acting as early warning signals of cell state changes [14,15]. Furthermore, eRNAs are functional players that help fold DNA (chromatin looping) [16] and release stalled polymerase [17], driving transcription. However, measuring eRNAs one by one is challenging because they are scarce and unstable, making them hard to use in standard analysis. We solve this by treating enhancers as coordinated groups (enhancer sets), using enrichment statistics to combine weak individual signals into strong regulatory readouts.

### No current method integrates enhancer-level evidence with gene-set enrichment

Existing regulatory-enrichment methods answer a different question than pathway corroboration. Gene-level enrichment lacks mechanistic context, and methods that infer gene regulatory networks (GRNs) from bulk RNA-seq are undercut by a basic limitation: mRNA abundance is an indirect, often lagging proxy for the protein-level activity that actually drives a pathway [18–21]. As a result, a transcription factor might drive a huge program without changing its own expression, making it invisible to standard network methods. While tools such as GREAT [22] and Lisa [23] have advanced the functional interpretation of regulatory regions, they operate on chromatin or TF binding data rather than integrating enhancer-level evidence with standard pathway enrichment output. More broadly, several established families interpret regulatory regions, yet none scores enhancer activity against pathway calls derived from the same experiment. Genomic region-set enrichment tools such as LOLA [24] and GIGGLE [25] take a pre-defined query set of regions—typically peaks called from a separate ChIP-seq or ATAC-seq assay—and test its spatial overlap against reference region-set databases; these tests are undirected and carry no expression or activity signal, so they cannot report whether a program is gained or lost in a given contrast. A related family—including count-based methods such as ChIP-Enrich [26] and Poly-Enrich [27], which incorporates chromatin long-range interactions—maps regions to genes (as GREAT does) and then applies conventional gene-set enrichment, using enhancers only as an intermediate to a gene list rather than as analytical units. Closest in spirit is chromVAR [28], which scores sets of peaks by chromatin-accessibility deviation; however, it requires accessibility data (scATAC-seq or ATAC-seq) rather than transcriptional output, estimates per-cell deviations for transcription-factor variation and clustering rather than a directional two-condition contrast, and is not designed to adjudicate pathway enrichment. Proximity-based approaches that ask whether a pathway’s regions lie near enhancer annotations [29] refine the interpretation of a gene-set result but introduce no independent measurement of enhancer activity. Critically, no existing method compiles an enhancer layer directly from enhancer RNA (eRNA) signal already present in the same transcriptome sequencing run, scores curated enhancer programs against a directional per-enhancer differential, and uses that orthogonal, condition-specific readout—obtained at no additional experimental cost—to corroborate gene-level pathway calls. This is the gap EnSEMBLE is built to fill.

### EnSEMBLE retains only pathways corroborated by an independent enhancer program

Here we introduce EnSEMBLE (Enhancer-Set Enrichment & Mechanism-Based Linked Evidence), a framework built on exactly that principle. EnSEMBLE pairs conventional gene-level GSEA with Enhancer-Program Enrichment Analysis (EPEA) over our curated enhancer-program libraries, then applies a deliberately strict logic: it collapses redundant pathway hits into concise process-level Themes and retains only those corroborated by a concordant, high-confidence enhancer program—a Helper. This dual-evidence requirement is the source of both EnSEMBLE’s confidence and its parsimony: it discards signals that the gene level alone would overcall, compressing hundreds of enrichments to a handful of evidence-backed claims (a reduction exceeding 97%) while making explicit which calls are trustworthy enough to act on. Because every step relies on fixed statistics and a transparent evidence classifier, each claim is traceable from raw signal to final verdict. Had such an enhancer-anchored readout been available for the *BCL2L14-ETV6* study, the mesenchymal program would have surfaced as a solid lead pathway at the outset rather than after a year of misdirection.

### The same enhancer libraries map non-coding GWAS variants to their target pathways

Finally, linking regulation to function is also critical in genetics. In Genome-Wide Association Studies (GWAS), most disease variants are in non-coding regions [30]. Traditional methods that link variants to the “nearest gene” might miss signals [31]; maps of DNA folding [32], and gene regulation [33] show enhancers often regulate distant targets. To address this, we include eORA (enhancer over-representation analysis), a tool that interprets GWAS variants with enhancer sets used in EnSEMBLE. By showing that our enhancer libraries can accurately map genetic traits to the correct tissues, we validate the resources used in our framework. We tested EnSEMBLE on GWAS Catalog datasets [34], showing it can recover expected biology and highlight interesting cases where regulation and expression disagree.

## Results

### EnSEMBLE corroborates gene-level pathway enrichment with enhancer-program evidence across four perturbation datasets

The principle behind EnSEMBLE is simple: a pathway call is trustworthy only when two independent layers of the same RNA-seq experiment agree on it. The first layer is the familiar one—gene-level enrichment (GSEA), which asks whether the genes of a pathway move together. The second layer reads the cell’s regulatory wiring directly: active enhancers transcribe short enhancer RNAs (eRNAs) that are captured in the same sequencing run, so the coordinated activity of an enhancer program can be scored alongside the genes it controls. EnSEMBLE keeps only those pathways whose gene-level signal is corroborated by a concordant enhancer program—what we term a Helper—and discards the rest.

To test this idea, we applied EnSEMBLE to four perturbation RNA-seq datasets spanning distinct cellular contexts and perturbation types (Table 1; three total RNA-seq datasets and one poly(A) mRNA-seq dataset). The perturbations include settings with well-characterized expected pathway shifts (EMT suppression after *SNAI1* knockout in MDA231; reduced translation and proliferation after rapamycin treatment in PANC1), a developmental transition from pluripotency to cortical neurons (iPSC-Neuron), and a setting where enhancer-level signal is attenuated by poly(A) selection (BT20 *BCL2L14-ETV6* Fusion).

**Table 1.**
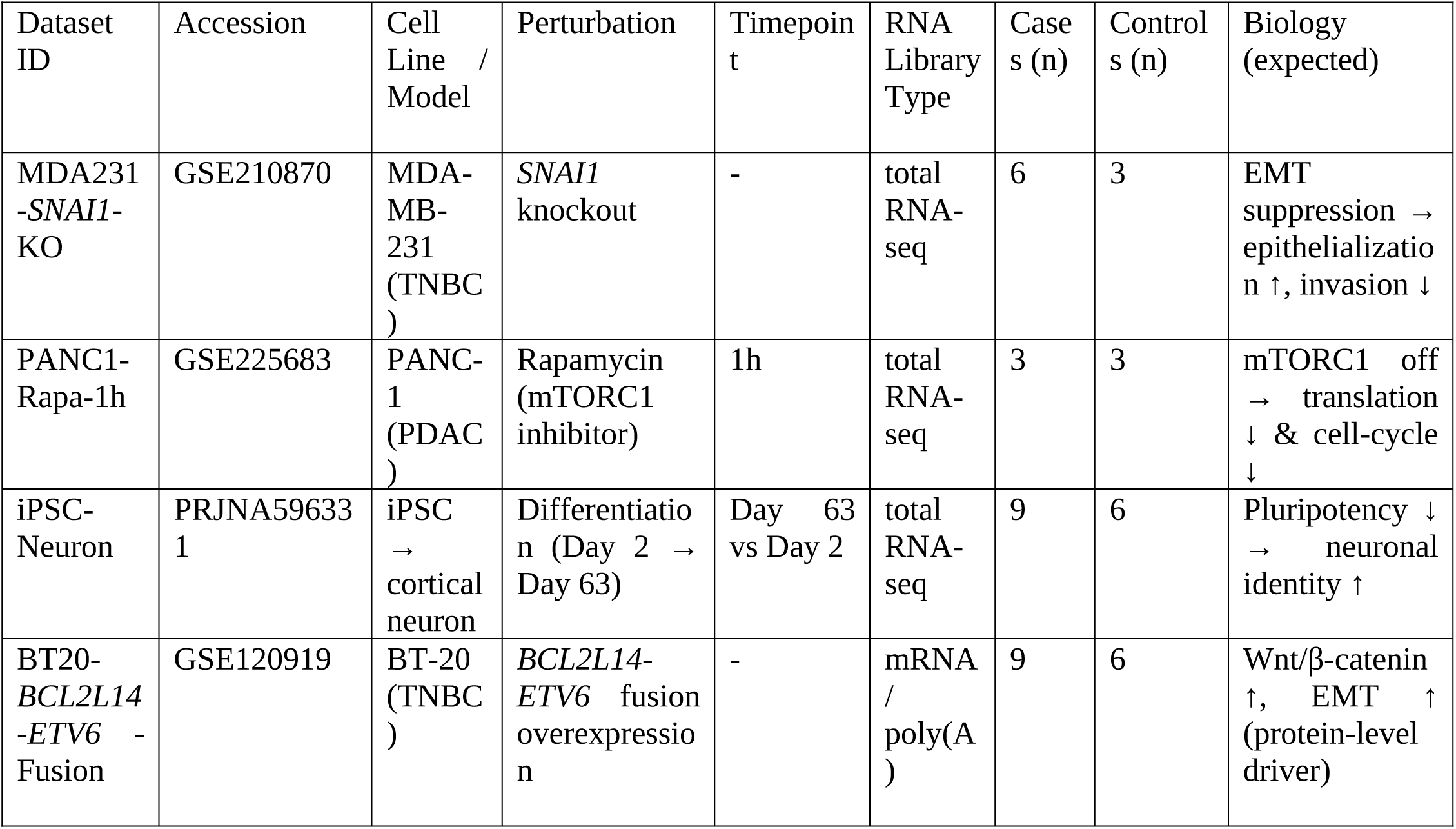
Validation datasets analyzed in this study.

| Dataset ID | Accession | Cell Line / Model | Perturbation | Timepoint | RNA Library Type | Cases (n) | Controls (n) | Biology (expected) |
| --- | --- | --- | --- | --- | --- | --- | --- | --- |
| MDA231-*SNAI1*-KO | GSE210870 | MDA-MB-231 (TNBC) | *SNAI1* knockout | - | total RNA-seq | 6 | 3 | EMT suppression → epithelialization ↑, invasion ↓ |
| PANC1-Rapa-1h | GSE225683 | PANC-1 (PDAC) | Rapamycin (mTORC1 inhibitor) | 1h | total RNA-seq | 3 | 3 | mTORC1 off → translation ↓ & cell-cycle ↓ |
| iPSC-Neuron | PRJNA596331 | iPSC → cortical neuron | Differentiation (Day 2 → Day 63) | Day 63 vs Day 2 | total RNA-seq | 9 | 6 | Pluripotency ↓ → neuronal identity ↑ |
| BT20-*BCL2L14-ETV6* - Fusion | GSE120919 | BT-20 (TNBC) | *BCL2L14-ETV6* fusion overexpression | - | mRNA / poly(A) | 9 | 6 | Wnt/β-catenin ↑, EMT ↑ (protein-level driver) |

The BT20 dataset derives from a TNBC cell line ectopically expressing *BCL2L14-ETV6* fusion variants, a class of *ETV6* structural rearrangements recently identified as highly specific to TNBC. These fusions act as dominant-negative alleles of wild-type *ETV6* and prime a partial epithelial-mesenchymal transition that enhances cell migration and invasion [11], providing a setting where enhancer-level and gene-level signals may capture downstream transcriptional consequences rather than the primary genetic lesion. Across datasets, EnSEMBLE integrates gene-level pathway enrichment with enhancer-program readouts and carries forward a compact set of helper-linked Themes for downstream evaluation.

### Curated enhancer-program libraries provide a regulatory-state readout for EPEA

Scoring the enhancer layer requires knowing which enhancers act together as a program. We therefore curated two complementary libraries of enhancer programs—sets of enhancers that share a regulatory identity (Fig. 1a). The first groups enhancers by the cell type in which they are active; the second groups them by the transcription-factor binding sites (TFBS) they contain. Before curation, the assembled collections comprised 945 enhancer sets across contributing catalogs, dominated by TFBS-defined programs (n=535, 56.6%) and CATlas cell-type programs (n=222, 23.5%) (Fig. S1a). Set sizes spanned orders of magnitude across sources (Fig. S1b), and a cosine-similarity network of the pre-filter cell-type programs showed substantial redundancy within and across broad family labels (Fig. S1c).

**Figure 1.**
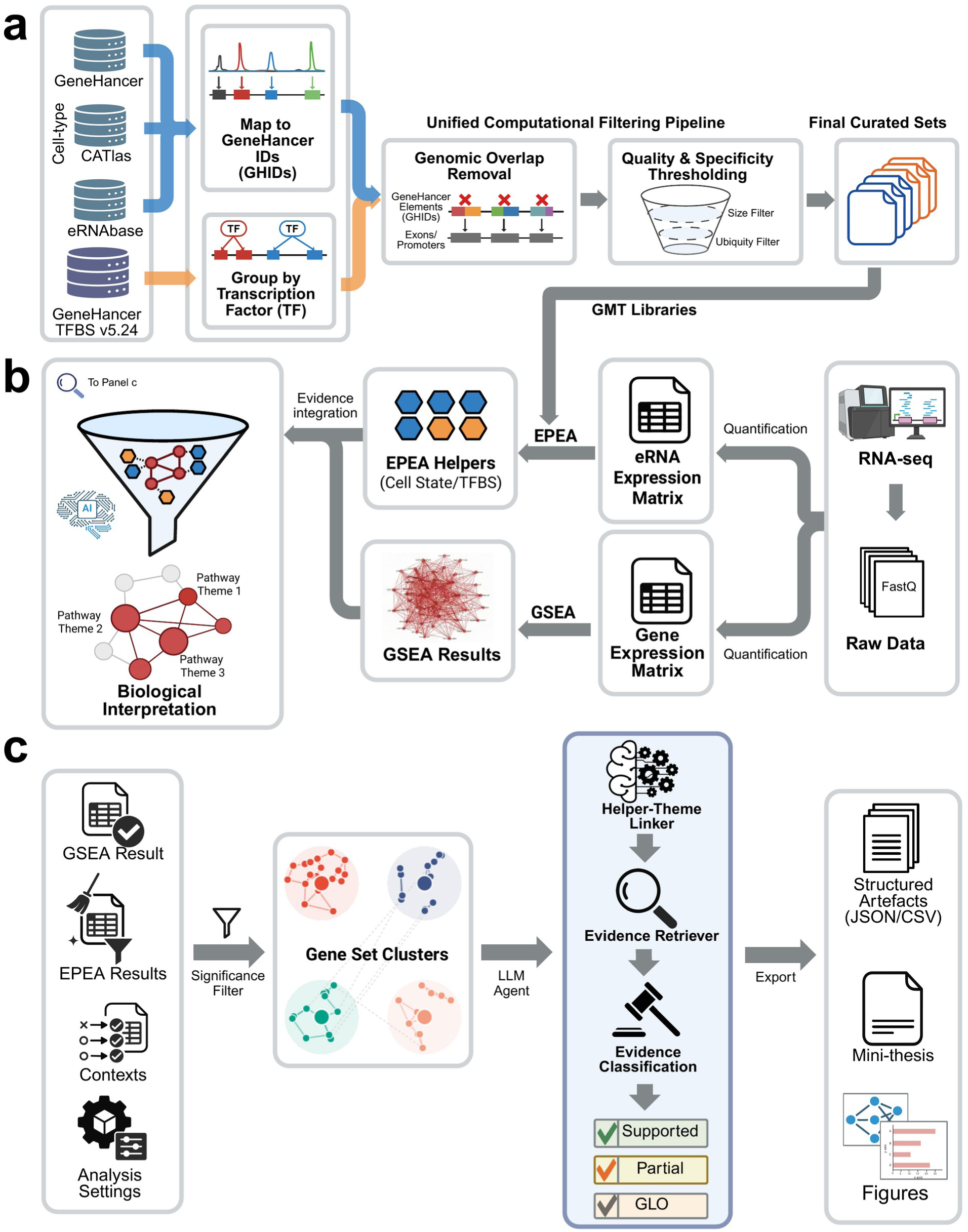
| EnSEMBLE overview. (a) EnSEMBLE curates two complementary enhancer-set libraries for EPEA: cell-type-specific enhancer sets (from GeneHancer, CATlas, and eRNAbase) and TFBS-defined enhancer sets (from GeneHancer TFBS). All elements are harmonized to GeneHancer enhancer IDs (GHIDs). A unified filtering pipeline removes exon/promoter-overlapping regions and applies size and ubiquity/specificity filters, yielding the final GMT libraries used throughout. (b) EnSEMBLE takes perturbation RNA-seq as input and derives gene-level and enhancer/eRNA-level differential signals. Gene-level pathway enrichment (GSEA) and enhancer-program enrichment (EPEA) are performed in parallel using curated reference libraries, producing pathway-scale Themes and enhancer-program Helpers. (c) Significant GSEA gene sets are consolidated into Themes by leading-edge Jaccard clustering. A structured evidence classifier then links Themes to Helpers and assigns verdict tiers (Supported, Partial, or Gene-level only) based on the specificity of the biological relationship. The classifier operates on precomputed summaries and is an optional interpretation aid (Methods). Outputs include structured artefacts (JSON/CSV), dataset-level narrative reports (mini-theses), and summary figures.

After standard library curation, both the number of retained programs and the distribution of set sizes were reshaped, with very small programs reduced and typical program granularity shifted toward more interpretable ranges (Fig. S2). All elements were harmonized to a unified enhancer identifier space (GeneHancer enhancer IDs; GHIDs) to enable consistent downstream enrichment and cross-library comparison; the resulting GMT libraries are provided as Supplementary Data and used throughout the EnSEMBLE analyses.

In downstream interpretation, we treat cell-type- and TFBS-labeled enrichments as regulatory-program readouts rather than direct evidence of cellular admixture or transcription-factor upregulation (Box 1). Unless otherwise stated, total RNA-seq datasets were analyzed using the full curated enhancer-program libraries, whereas the BT-20 mRNA-seq analysis used a sensitivity-oriented restricted collection (Methods; Table S1).

#### Box 1

**| How to read EnSEMBLE output**

A Helper is an enhancer program that shows significant coordinated activity in the same comparison as the gene-level result (by enhancer-set enrichment analysis, EPEA). It provides an independent, regulatory line of evidence for interpreting a gene-level Theme. The direction of a Helper is read from its normalized enrichment score (NES): NES greater than 0 means the program is more active in the case (e.g., treated or perturbed) condition, and NES less than 0 means it is less active. Helper names describe regulatory identity, not cell content: a cell-type-named Helper (e.g., “fibroblast”) means the responding enhancers resemble that cell type’s regulatory program, not that fibroblasts are present in the sample; a transcription-factor-binding-site (TFBS) Helper means the responding enhancers are enriched for that factor’s binding sites, not that the factor’s own mRNA is up.

Each Theme is then assigned one of three verdict tiers that grade how strongly the enhancer evidence backs it. Supported means a Helper’s regulatory identity defines the Theme—for example, a glutamatergic-neuron Helper linked to a synaptic-transmission Theme. Partial means the Theme reflects a characteristic but non-defining activity of the Helper’s program. Gene-level only means the pathway was significant at the gene level but no matching enhancer program was found, so the call rests on gene evidence alone. Before a Helper is trusted, its reliability is checked two ways: it must stay significant when individual replicates are dropped (leave-one-replicate-out, LORO, stability) and must outperform random enhancer sets of the same size (set-identity shuffles) (Fig. 4).

### EnSEMBLE integrates gene-level and enhancer-level enrichment into verdict-tagged claims

As summarized in Fig. 1b-c, EnSEMBLE starts from perturbation RNA-seq and jointly analyzes gene-level pathway enrichment and enhancer-program enrichment to connect process-level changes to regulatory context. In brief, gene-level GSEA identifies which pathways changed, while enhancer-program enrichment (EPEA) asks which enhancer programs shifted in the same comparison. Significant enhancer programs become Helpers—compact, interpretable readouts of regulatory state. EnSEMBLE then collapses the redundant pathway hits into process-level Themes and reports only those Themes a Helper corroborates. Across datasets, enhancer-level differential activity was detectable and varied by assay type, providing the substrate for EPEA and motivating library configurations tailored to total RNA-seq versus poly(A) mRNA-seq inputs (Fig. S3; Methods).

### Two-stage consolidation reduces reported signatures by more than 97%

A standard GSEA run returns far more significant gene sets than anyone can act on. EnSEMBLE tames this in two steps: it first groups redundant gene sets into process-level Themes, then keeps only the Themes an enhancer program supports (Methods; Fig. 2a). The compression is dramatic. Using gene-level GSEA across four MSigDB collections (KEGG, GOBP, REACTOME, HALLMARK), we observed 266, 402, 870, and 775 significant gene sets (FDR < 0.05) in BT20 *ETV6*-Fusion, MDA231 *SNAI1*-KO, PANC1 Rapa-1h, and iPSC-Neuron, respectively. Data-driven clustering condensed these into 32, 26, 37, and 37 Themes, and the evidence classifier identified 3, 10, 6, and 18 final claims (Supported or Partial), corresponding to a 97.5–99.3% reduction in the number of reported signatures (Fig. 2a,c). Crucially, this compression is not mere pruning: as shown below, the handful of claims that survive are the biologically expected ones, so EnSEMBLE discards an order of magnitude of redundancy without discarding the signal.

**Figure 2.**
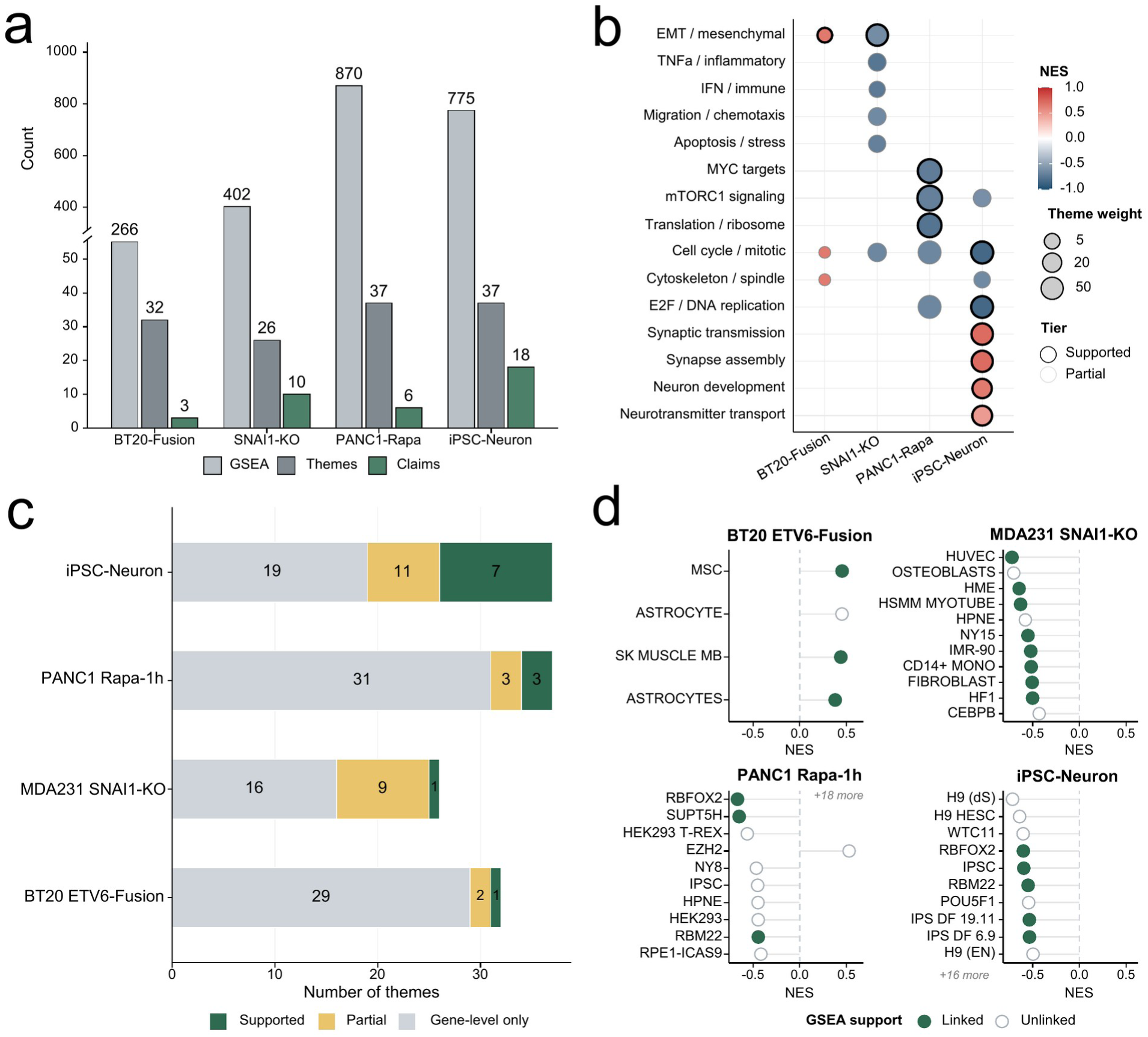
| EnSEMBLE compresses GSEA hits into helper-supported claims (a-d). (a) Number of significant gene sets from gene-level GSEA (FDR < 0.05), process-level Themes after clustering, and final claims (Supported or Partial verdicts) per dataset. Numbers above bars indicate counts. (b) Bubble heatmap summarizing helper-supported Themes across the four datasets. Each row represents a broad biology category; color denotes Theme effect direction (NES; red = up in case, blue = down in case); bubble size indicates evidence weight; solid border denotes Supported verdict, dashed border denotes Partial. (c) Verdict composition per dataset shown as stacked bars. Green: Supported; gold: Partial; gray: Gene-level only. (d) Overview of selected EPEA helpers per dataset. Each point shows one helper’s NES from EPEA. Filled circles indicate helpers linked to at least one Supported or Partial claim; open circles indicate helpers not linked to any final claim. Helpers are ranked by absolute NES within each dataset.

To support traceability, EnSEMBLE exports Theme membership and Theme-level summaries together with a structured claim table linking Themes to Helpers and recording verdict tiers (Supplementary Tables S5–S6; Supplementary Data).

### Helper-linked Themes recover expected biology and resolve regulatory mechanism

The test of any prioritization method is whether the claims it keeps are the biologically right ones. Across all four datasets, the Helper-supported Themes recovered the expected biology—and, in several cases, named a regulatory mechanism that gene-level analysis alone had missed (Fig. 2b–d).

#### I. *SNAI1* knockout collapses a coordinated mesenchymal program

*SNAI1* is a master regulator of EMT, so knocking it out should collapse the mesenchymal program—and that is exactly the top claim EnSEMBLE returned. In MDA231 *SNAI1*-KO, the EMT Theme received the strongest evidence weight of any claim across all datasets (w = 31.3) and was downregulated, supported by three independent mesenchymal enhancer programs (HSMM myotube, fibroblast, and hair follicle HF1; Fig. 3b). This convergence from distinct reference programs strengthens the conclusion that *SNAI1* knockout dismantles a coordinated mesenchymal regulatory state, consistent with *SNAI1*’s established role in maintaining the mesenchymal program whose loss drives a partial mesenchymal-to-epithelial transition [35,36].

**Figure 3.**
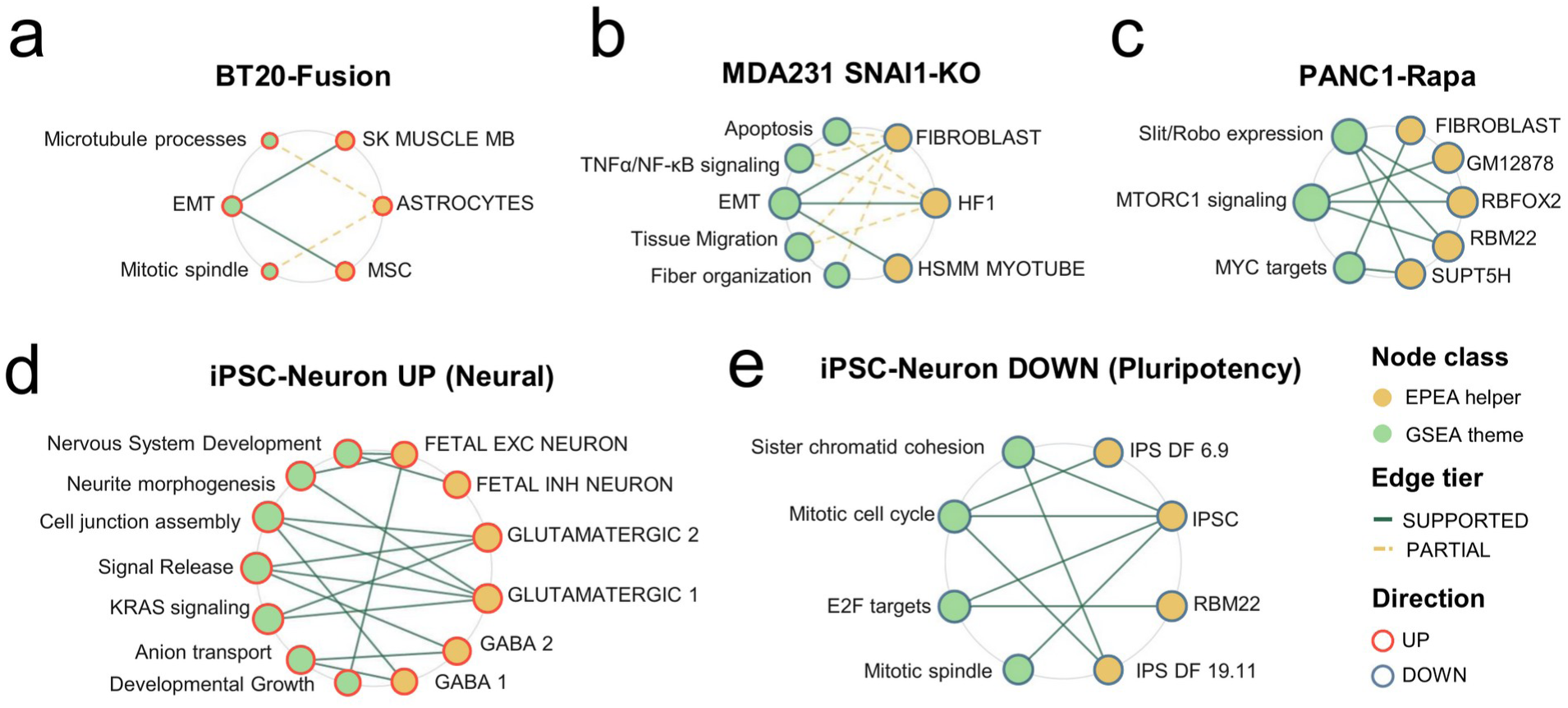
| Bipartite networks link Helpers to Themes across perturbations. **(a-e).** Network diagrams visualizing evidence links between Helpers (green nodes, right) and biological Themes (blue nodes, left) for each dataset. Solid green edges indicate Supported verdicts; dashed gold edges indicate Partial verdicts. Only Themes with at least one Supported or Partial verdict are shown; Gene-level only Themes are omitted. (a) BT20 *ETV6*-Fusion. (b) MDA231 *SNAI1*-KO. (c) PANC1-Rapa. (d, e) iPSC-Neuron, shown as two direction-specific sub-panels to visualize the bilateral regulatory convergence unique to this dataset: (d) UP direction (neural maturation Themes linked to neuron-subtype-specific helpers) and (e) DOWN direction (proliferative/pluripotency Themes linked to iPSC-type helpers).

Ten further Themes received Partial verdicts. These included a coherent set of downregulated immune and inflammatory programs, namely TNFα/NF-κB signaling, IFN-γ response, immune/allograft-rejection, and cell chemotaxis, each linked to shared CD14+ monocyte and fibroblast enhancer programs. Because EnSEMBLE treats cell-type-labeled helpers as regulatory-program proxies rather than evidence of cellular admixture (Box 1), this pattern indicates that the collapsing mesenchymal program is accompanied by coordinated loss of inflammatory enhancer activity, an EMT-coupled inflammatory state consistent with the reciprocal *SNAI1* and NF-κB regulation described previously [37]. The remaining Partial Themes captured migration, actin and extracellular-matrix organization, and apoptotic programs, all characteristic of, but not specific to, the mesenchymal state. The downregulation of the EMT and TGFβ-associated programs is directly concordant with the source study, which demonstrated that *SNAI1* knockout induces a partial mesenchymal-to-epithelial transition with attenuated TGFβ signaling [35].

#### II. The rapamycin response reveals an mTOR–MYC–SPT5 elongation axis

The PANC1 case shows the payoff of the enhancer layer most clearly: it surfaced a mechanistic link that the gene list alone left invisible. One hour after rapamycin treatment, the Themes receiving the highest evidence weights were mTORC1 signaling, MYC targets, and a ribosome-biogenesis program (labeled by its representative gene set; Methods), all downregulated and all Supported (w up to 76.7; Fig. 3c). The mTORC1 signaling Theme was linked to the *RBFOX2*, *RBM22*, and GM12878 enhancer programs, each of which carries the HALLMARK mTORC1 signaling gene set as its top annotation. The MYC targets Theme was linked to *SUPT5H* and a fibroblast program, with *SUPT5H* carrying the HALLMARK MYC targets gene set as its top annotation. This *SUPT5H* link is consistent with the predicted mTOR to MYC to SPT5 elongation axis: mTORC1 promotes *MYC* translation [38], *MYC* promotes transcriptional pause release via P-TEFb [39], and *MYC* directly recruits SPT5 (*SUPT5H*), a component of the DSIF elongation complex, to Pol II to enable processive elongation [40].

Because the HALLMARK mTORC1 signaling gene set captures transcriptional targets downstream of mTOR activity rather than the phospho-signaling components themselves [41], the Supported verdict indicates that these transcriptional consequences are accompanied by concordant enhancer-level changes within one hour of rapamycin treatment, a timescale consistent with rapid eRNA dynamics in other signaling contexts [14].

The source study characterized rapamycin’s effect at the translational level using ribosome profiling, where mTOR inhibition triggers feedback activation of a subset of mRNAs [42]. EnSEMBLE instead operates on steady-state transcript abundance and enhancer activity, and recovers the coordinated downregulation of mTORC1 and MYC target transcripts together with their supporting enhancer programs. The two readouts are complementary: transcriptional and enhancer-level repression of the mTORC1/MYC program in our analysis accompanies the translational feedback described in the source study.

#### III. Dual-evidence filtering favors enhancer-supported EMT over dominant proliferative signal

BT20-BCL2L14-*ETV6*-Fusion illustrates the opposite and equally important move: the dual-evidence requirement demoting a statistically dominant but mechanistically unsupported signal. Here gene-level enrichment was dominated by proliferative signatures, with E2F targets (NES = +0.88), G2/M checkpoint, and DNA-replication and repair Themes ranking as the most significant gene-level signals. EnSEMBLE assigned all of these Gene-level only verdicts, because none of the recovered enhancer programs had a regulatory identity defined by cell-cycle control, so the dual-evidence requirement was not met. In contrast, the EMT Theme, although weaker at the gene level, was the only Theme to receive concordant enhancer support, linked to two mesenchymal-lineage programs (mesenchymal stem cell and skeletal muscle myoblast; EPEA q ≈ 0.045), and was promoted to Supported. This prioritization, which demotes the statistically dominant but regulatory-context-poor proliferative signal while retaining the enhancer-supported mesenchymal program, is concordant with the functional characterization of these fusions, which enhance cell migration and invasion but do not alter cell viability or cell-cycle progression in BT20 cells [11].

Among the two mesenchymal-lineage programs supporting this verdict, the skeletal muscle myoblast program was specific under the set-identity shuffle (empirical p = 0.013), whereas the mesenchymal stem cell program yielded too few valid size-matched nulls for a reliable shuffle test (3 valid nulls; Supplementary Table S4). The Supported verdict therefore rests on the concordant gene-level and EPEA enrichment together with the myoblast shuffle specificity and leave-one-sample-out stability (Table S3). The attenuated overall enhancer signal in this dataset is consistent with reduced eRNA capture under poly(A) selection (Methods).

#### IV. A complete developmental transition yields the most helper-linked claims

A complete developmental transition gave EnSEMBLE the most to work with—and showed that the approach generalizes well beyond cancer cell lines. In the iPSC-to-neuron differentiation, EnSEMBLE identified the most helper-linked claims of any dataset (11 Supported, 7 Partial), with both up- and downregulated directions showing regulatory convergence (Fig. 3d). Upregulated neural Themes (synaptic transmission, cell junction assembly, neuron development) were Supported by neuron-subtype-specific enhancer programs (glutamatergic, GABAergic, fetal excitatory and inhibitory neuron programs), independently recovering the neuronal identity validated by electrophysiology and immunostaining in the source study [43]. Downregulated proliferative Themes (E2F targets, mitotic cell cycle, sister chromatid cohesion) were Supported by iPSC-type enhancer programs, capturing the established coupling between cell-cycle exit and pluripotency loss [44].

The higher number of Supported claims in this dataset reflects the strength of a complete cell-identity transition (Day 2 to Day 63), well-matched cell-type enhancer libraries, and the larger sample size (15 samples, 5 donors), rather than methodological bias. This non-cancer developmental dataset also demonstrates that EnSEMBLE generalizes beyond in vitro cancer perturbations.

### Verdict tiers filter enrichments lacking enhancer support

Finally, the verdict framework served as an automated filter, downgrading enrichments that lacked concordant enhancer support. Critically, these downgrades are reported as explicit statements of insufficient evidence rather than silently dropped: a Gene-level only verdict declares that regulatory corroboration was not found, not that the pathway is absent—turning what would otherwise be an invisible false negative into an honest, auditable uncertainty. A Hedgehog-signaling enrichment in PANC1 was classified as Gene-level only because its leading-edge was dominated by housekeeping modules without corresponding enhancer-program support. Similarly, IFN-α signatures in iPSC-Neuron were not promoted to Supported, consistent with the absence of interferon-related enhancer programs in a clean in vitro differentiation system (Table S5).

### Claim-linked Helpers are stable, specific, and condition-dependent

A claim is only as trustworthy as the Helper behind it, so we subjected every claim-linked Helper to three independent tests: does it survive dropping samples, is it more than a generic large set, and does it depend on the actual case/control contrast? First, elbow-based selection reduced the significant EPEA hits to a compact subset of enhancer-program readouts for each total RNA-seq perturbation (Table S2; Methods). To focus on the helpers most relevant to downstream interpretation, the main-text robustness panels (Fig. 4b–d) report only the subset of helpers linked to final Supported or Partial claims (20 helpers across four datasets); complete per-helper results for all selected helpers are provided in Supplementary Tables S3–S4.

**Figure 4.**
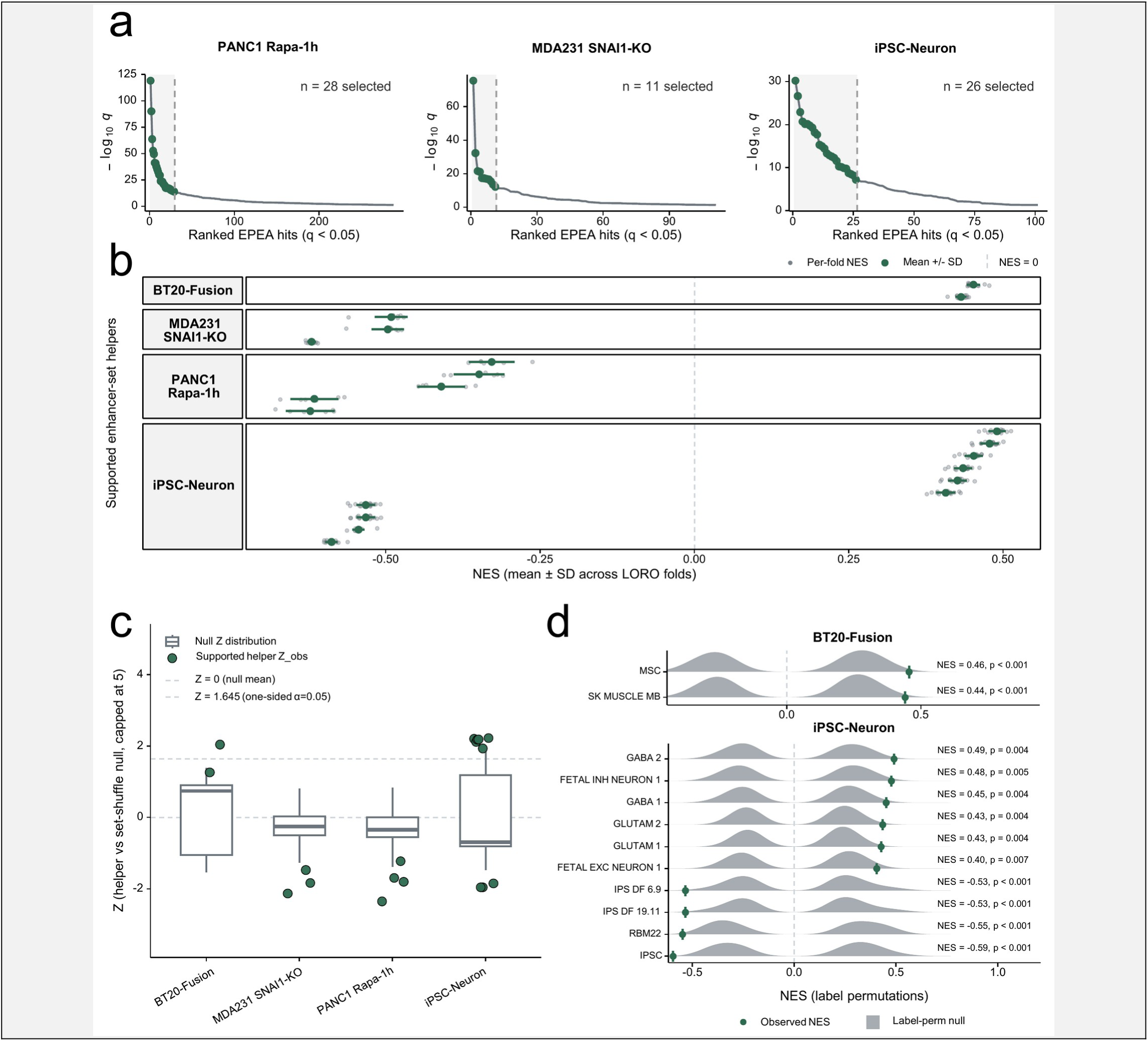
| Helper selection, stability, and specificity (a-d). (a) Kneedle-style elbow selection applied to ranked significant EPEA hits (q < 0.05) pooled across cell-type and TFBS-defined enhancer-set libraries for the three total RNA-seq datasets (MDA231 *SNAI1*-KO, PANC1-Rapa, iPSC-Neuron). Green points indicate selected helpers; gray points indicate other significant hits below the elbow cutoff. For BT20 (poly(A) mRNA-seq), all significant hits were retained without elbow selection (Methods). (b) Leave-one-replicate-out (LORO) stability of selected helpers, summarized as mean NES with standard deviation across folds. Larger green points indicate helpers linked to Supported or Partial claims; smaller gray points indicate unlinked helpers. (c) Set-identity shuffle specificity for selected helpers across the four datasets. Each helper is compared to a size-matched null distribution generated by randomizing set membership; the y-axis shows the standardized specificity score (Z_dir) relative to each helper’s own null. The dashed line marks Z_dir = 1.645 (one-sided p = 0.05). (d) Label permutation analysis for BT20 *BCL2L14-ETV6*-Fusion (P = 1,000 permutations) and iPSC-Neuron (P = 1,000 permutations). Gray density shows the distribution of helper NES under randomized case/control labels; green diamonds indicate the observed NES. Empirical p-values are reported at permutation resolution.

These claim-linked helpers were highly stable under leave-one-replicate-out reanalysis, showing consistent enrichment directions and magnitudes across folds in all four datasets, including the iPSC-Neuron dataset where each of 15 LORO folds removed a single sample under a donor-blocked design (Fig. 4b; Table S3). Helpers were also generally well separated from size-matched set-identity shuffle nulls, indicating that observed enrichments depend on specific enhancer-program composition rather than generic properties of similarly sized sets (Fig. 4c; Table S4).

To further test whether helper signals could arise under arbitrary case/control assignments, we performed label permutation analyses for BT20-*BCL2L14-ETV6*-Fusion (P = 1,000) and iPSC-Neuron (P = 1,000; unrestricted global permutation with DESeq2 donor blocking refit per permutation). In both datasets, observed helper NES values fell outside the permutation null distributions, confirming that helper enrichments are condition-dependent rather than artifacts of sample structure (Fig. 4d). For the BT-20 poly(A) mRNA-seq dataset, we used the mRNA sensitivity configuration to accommodate attenuated enhancer-level signal (Methods; Table S1).

### eORA recovers cell-type–selective enhancer programs from GWAS associations

If our enhancer libraries genuinely capture cell-type regulatory identity, they should be able to recover the right tissue from disease genetics alone—a test that has nothing to do with the perturbation experiments. We put them to exactly this test. Most disease-associated variants fall in non-coding enhancers, so a trait’s causal cell type should be encoded in which enhancer programs its variants land in. We applied eORA (enhancer over-representation analysis) to 1,575 QC-passed GWAS Catalog traits to ask whether the curated enhancer-program library yields structured, cell-type–selective signals in this orthogonal setting (Fig. 5a–b; Supplementary Table S9). Trait-by-program enrichments showed clear family-level structure rather than diffuse associations (Fig. 5b).

**Figure 5.**
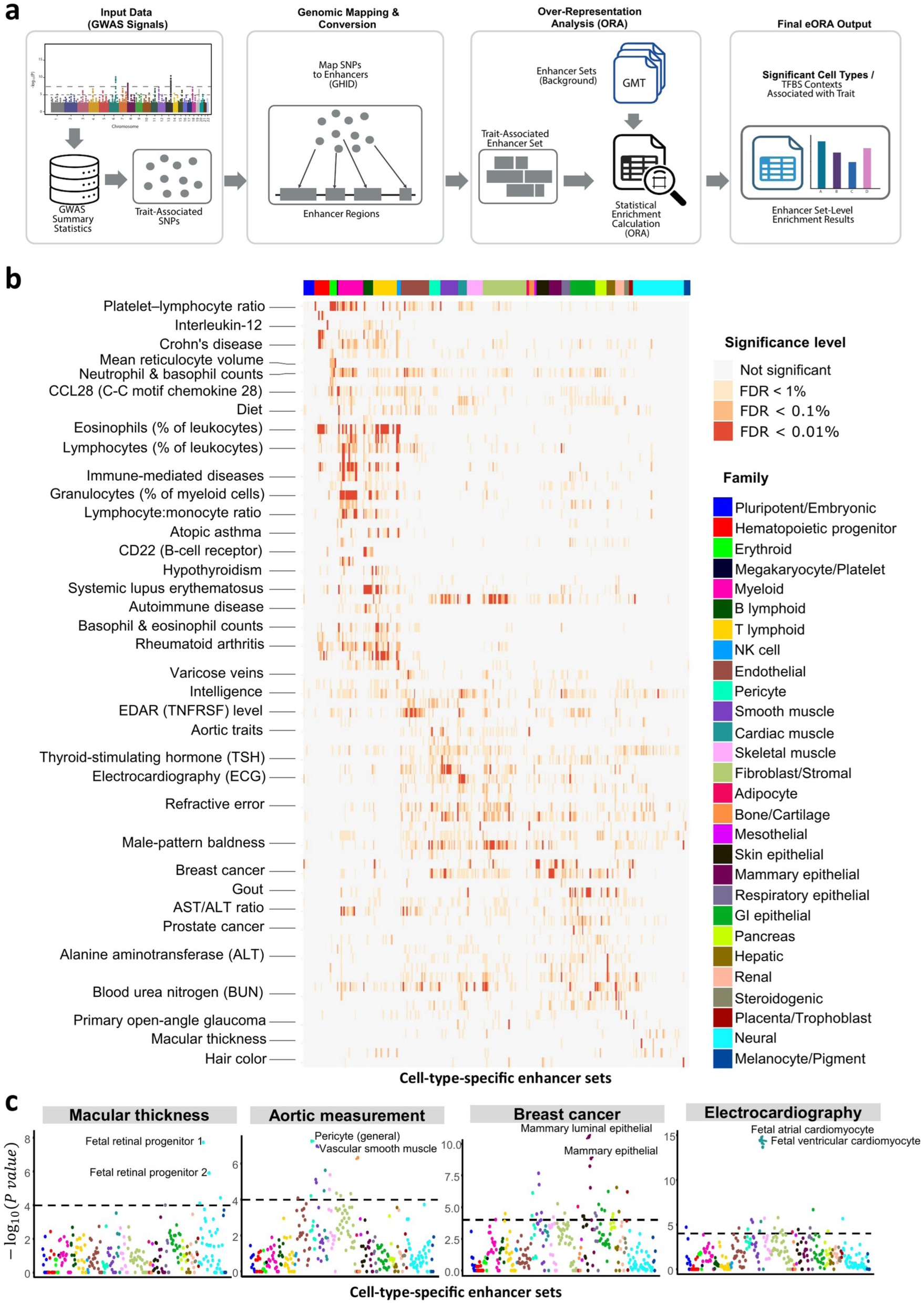
| eORA connects GWAS loci to cell-type enhancer programs (a-c). (a) eORA workflow. For each GWAS trait, lead SNPs are mapped to GeneHancer enhancers (GHIDs) to define a trait-associated enhancer set, followed by over-representation analysis (hypergeometric test) against curated cell-type enhancer programs. (b) Overview heatmap of selected GWAS Catalog traits versus cell-type enhancer programs, with significance binned by trait-wise FDR (Benjamini-Hochberg correction within each trait). Enhancer-set family labels are used for visualization only and were not used in statistical testing. (c) Four representative traits shown as enhancer-program scans across all tested cell-type programs (y-axis: -log10(p)). The dashed line indicates the Bonferroni display threshold (0.05/489, where 489 is the number of tested cell-type programs). eORA results are based on cell-type enhancer programs only; TFBS-defined libraries were not included. Input coverage, mapping scale, and size-matched null calibration diagnostics are summarized in Fig. S5, and full trait-by-program results are provided in Supplementary Table S9.

In focused trait scans, the strongest signals were highly selective and biologically concordant: breast carcinoma was dominated by mammary epithelial enhancer programs (luminal/basal/myoepithelial), electrocardiography by cardiomyocyte programs, aortic measurement by vascular wall–associated programs (pericyte and vascular smooth muscle), and macula-related traits by retinal programs (retinal progenitor and pigment-related sets) (Fig. 5c). Notably, these selective peaks were observed despite small trait-associated enhancer set sizes (median mapped GHIDs per trait = 11; Fig. S5a) and were consistent with conservative null-calibration diagnostics (Fig. S5c), supporting that the observed enrichments reflect specific enhancer-program identity rather than artifacts of mapping scale or p-value inflation.

## Discussion

### A verification step that turns pathway lists into adjudicated claims

Pathway enrichment remains a standard tool for transcriptomics, yet it creates an interpretation bottleneck by producing long lists of redundant gene sets that encourage subjective selection. EnSEMBLE resolves this not by creating a new statistic, but by enforcing a verification step: pairing gene-level pathways with orthogonal enhancer-program readouts. The key enabling insight is that this second evidence layer comes for free. Active enhancers transcribe enhancer RNAs (eRNAs) that are already present in standard RNA-seq, so the regulatory state of a pathway can be scored from the very same data that produced the gene-level signal—no new assay, cost, or sample is required. This turns a near-universal but underused signal into a routine corroboration layer for pathway analysis. By consolidating redundant signals into process-level Themes and anchoring them to high-confidence Helpers, EnSEMBLE reduces reported signatures by >97%, shifting the analytical focus from enumeration of possibilities to adjudication of evidence.

### The Helper concept: enhancer activity as orthogonal evidence

A central concept is the Helper: a significant enhancer-program enrichment that serves as a proxy readout of coordinated regulatory activity. Helpers should not be interpreted as cell-type contamination or TF mRNA upregulation (Box 1). Each Theme is classified as Supported, Partial, or Gene-level only based on the specificity of its relationship to direction-matched Helpers, providing a graduated evidence framework rather than a binary filter.

### From describing expression to resolving regulatory mechanism

The practical value of EnSEMBLE lies not in compression per se, but in the qualitative shift from describing what genes changed to identifying what regulatory programs are driving those changes. In MDA231 *SNAI1*-KO, three independent mesenchymal enhancer programs (myotube, fibroblast, hair follicle) converged on the EMT Theme, pointing specifically toward mesenchymal enhancer state as the regulatory substrate being dismantled, a level of mechanistic resolution that standard GSEA cannot provide. In PANC1 Rapa-1h, the helper layer surfaced *SUPT5H* (SPT5), connecting mTOR inhibition to transcriptional elongation machinery via the predicted MYC-SPT5 axis [39,40]; this connection was invisible in the gene-level GSEA output alone. In iPSC-Neuron, neuron-subtype-specific helpers (glutamatergic, GABAergic, fetal excitatory) independently recovered the cellular identity of the differentiation from enhancer signals, demonstrating that helper-linked claims carry information about the regulatory identity of the transition, not merely its pathway composition. These examples illustrate how helper identity transforms pathway analysis from a descriptive exercise into a hypothesis-generating platform that can direct follow-up experiments toward specific regulatory mechanisms.

### Themes preserve conflicting evidence as explicit hypotheses

Across the perturbations analyzed, Themes recovered expected biology while also representing timing- or layer-dependent mismatches as explicit hypotheses rather than forcing a single narrative. For example, acute inhibition of mTORC1 suppressed translation-associated programs with concordant enhancer support, while compensatory PI3K-AKT signaling signatures, which the original study attributed to translational feedback [42], lacked enhancer-level backing in the transcriptional data. This discipline preserves a compact set of claims for communication while retaining a traceable record of supporting and conflicting evidence that can be used to prioritize validation.

### Relationship to existing regulatory-enrichment tools

Existing tools for regulatory enrichment analysis address related but distinct questions. GREAT [22] performs ontology enrichment on genomic regions; Lisa [23] infers transcriptional regulators from chromatin models; ChEA3 [45] ranks transcription factors by multi-library integration; and i-cisTarget [46] identifies enriched motifs and experimental tracks in gene sets. Genomic region-set enrichment tools (e.g., LOLA, GIGGLE) test spatial overlap of a pre-called region set against reference sets but are undirected and require a separate chromatin assay; chromVAR scores peak sets from chromatin-accessibility deviation rather than eRNA transcription and is oriented toward transcription-factor variation rather than pathway corroboration. None of these specifically layers enhancer-program evidence over GSEA pathway output as a second-tier prioritization step. EnSEMBLE fills this gap by operating downstream of standard pathway analysis, adding regulatory context to existing GSEA results rather than replacing them.

### eORA extends the enhancer libraries to disease genetics

We positioned eORA as a complementary application that reuses the same enhancer-program resources in a GWAS setting. EPEA addresses rank-based perturbation questions, specifically whether a program’s enhancers shift coherently along a differential signal, whereas eORA addresses list-based GWAS questions, specifically whether a trait’s mapped enhancers are over-represented in specific cell-type programs. The ability of eORA to recover known cell-type contexts from GWAS summary statistics serves as an independent validation of the underlying enhancer libraries, reinforcing confidence in the EPEA helpers detected in perturbation data.

### An agentic, fully reproducible discovery framework

Beyond the enhancer layer, EnSEMBLE reframes pathway interpretation as an agentic discovery task: a language agent reasons over precomputed, statistically grounded evidence to adjudicate claims, rather than a human scanning hundreds of enrichments by hand. Crucially, the agent is constrained to synthesis, not statistics. EnSEMBLE separates statistical inference from narrative synthesis. All differential analyses and enrichment statistics are computed deterministically; a structured classifier is used only to organize precomputed summaries into verdict-tagged claims under fixed output constraints. It serves as an interpretation aid that compensates for the practical difficulty of manually evaluating hundreds of Theme-Helper pairs against broad biological knowledge, not as an analytical engine. Users confident in their domain expertise may bypass the classifier entirely and interpret the upstream GSEA, EPEA, and Theme outputs directly. Reproducibility is supported by temperature-zero inference, deterministic validation rules, and a three-run consensus policy, following recent precedents for constrained LLM use in genomics [47,48]. All locked parameters, classifier prompts, and per-dataset audit artefacts are publicly available.

### Limitations and future directions

The conditions under which EnSEMBLE performs best also define its current scope. Because the enhancer layer scales with the magnitude of regulatory change, it is most powerful when enhancer profiles shift broadly, as during cell-state transitions, and less sensitive to subtle shifts. Sensitivity is highest with total RNA-seq, which captures eRNAs richly. Importantly, EnSEMBLE also works on poly(A) mRNA-seq: even with attenuated eRNA capture, it still detects large-scale enhancer reprogramming, extending the method to the vast archive of existing mRNA-seq data. Coverage is likewise shaped by the reference enhancer catalogs from which Helpers are drawn, so a missing Supported verdict signals an opportunity for library expansion rather than confirmed absence, and coverage will broaden as these catalogs grow. Two further design choices define the inputs: EnSEMBLE uses a user-supplied template to anchor the contrast, and eORA enrichments—shaped by LD structure and annotation coverage—are best read as cell-type context candidates. Demonstrated here across four cancer and developmental datasets, EnSEMBLE offers a foundation that future work can extend to new tissues, modalities, and organisms and benchmark against orthogonal measurements such as chromatin accessibility, transcription-factor occupancy, and direct enhancer perturbation.

## Methods

### Overview and notation

EnSEMBLE integrates gene-level pathway enrichment with enhancer-program enrichment to derive a compact set of evidence-tagged claims from perturbation RNA-seq. Briefly, we perform gene-level GSEA and enhancer-program enrichment analysis (EPEA), select enhancer-program readouts (Helpers), consolidate GSEA hits into Themes, and generate helper-Theme claims with a rule-constrained LLM agent then assembles traceable, structured outputs and adjudicates verdicts.

We use the following terms throughout: *GHID*, a harmonized enhancer identifier from GeneHancer; *enhancer set*, a curated set of enhancers representing a regulatory program; *Helper*, a selected significant EPEA hit used as a high-confidence program readout; *Theme*, a consolidated process-level summary of related GSEA gene sets; and *claim*, a helper-linked Theme with a verdict label (Supported / Partial / Gene-level only). We define final claims as those with verdict Supported or Partial; Gene-level only claims are retained for traceability.

### Perturbation RNA-seq datasets

We evaluated EnSEMBLE on four perturbation RNA-seq datasets (three total RNA-seq; one mRNA/poly(A) RNA-seq). Dataset identifiers, biological systems, perturbations, time points, replicate counts, accession identifiers, and case/control sample numbers are provided in Table 1.

For the three public datasets [35,42,43], we downloaded RNA-seq FASTQ files from SRA and processed all samples with our quantification workflow to generate a gene-by-sample matrix and a GHID-by-sample enhancer/eRNA matrix. Sample annotations and group labels were derived from repository metadata (Table 1).

Samples were included if they matched the Table 1 case/control definitions and had complete metadata. Uniform QC filters were applied across datasets with thresholds reported in Supplementary Table S1. Differential analyses used the unified contrast specification and produced gene- and GHID-level differential statistics for downstream GSEA and EPEA.

### Curated enhancer-set libraries enable regulatory-program readouts

To support enhancer-level interpretation, we curated two complementary enhancer set libraries for EPEA: cell-type enhancer sets and TFBS-defined enhancer sets (Fig. 1a). Cell-type programs were compiled from GeneHancer, CATlas, and eRNAbase cell-type collections, while TFBS-defined programs were derived from GeneHancer TFBS annotations by grouping enhancers by transcription factor. All elements were harmonized to a unified enhancer identifier space (GHIDs), enabling consistent downstream enrichment and cross-library comparison. Library composition and size characteristics are summarized in Fig. S1, and the effect of standard library curation on library complexity is summarized in Fig. S2; the resulting GMT libraries are provided as Supplementary Data and used throughout the EnSEMBLE analyses.

In downstream interpretation, we treat cell-type- and TFBS-labeled enrichments as regulatory-program readouts rather than direct evidence of cellular admixture or transcription-factor upregulation (Box 1). Unless otherwise stated, total RNA-seq datasets were analyzed using the full curated enhancer-program libraries, whereas the BT-20 mRNA-seq analysis used a sensitivity-oriented restricted collection (Methods; Table S1).

### Read processing and expression quantification

For datasets with raw reads, we processed FASTQ files with a single quantification workflow to generate (i) a gene-by-sample matrix and (ii) a GHID-by-sample matrix for enhancer/eRNA signal. Reads were aligned to the human reference genome (GRCh38 primary assembly) with Rsubread, retaining uniquely mapped reads, and quantified with featureCounts against GENCODE v47 (basic) gene annotations and GeneHancer v5.24 (hg38) enhancer intervals (BED) for enhancer/eRNA counting. Per-sample count tables were merged into expression matrices; features with zero counts across all samples were removed. The quantification workflow is implemented as open-source R code and available on GitHub.

Sample metadata were parsed from repository (public datasets) or internal (BT-20) annotations and aligned to the columns of the expression matrices by sample identifier. Only samples present in both the metadata table and the corresponding expression matrix were retained.

Prior to differential testing, we filtered low-expression features under the specified case-control design (primary expression filter). For enhancer/eRNA matrices, we additionally restricted to GeneHancer enhancers after removing entries annotated as promoters and those overlapping GENCODE exon annotations (consistent genome build).

Differential expression for both genes and GHIDs was performed with DESeq2 on filtered raw counts. Counts were rounded to integers (default) and features with very low total counts were removed (row sum ≤ 10 by default) as an input safeguard for DESeq2.

For downstream enrichment, we converted DESeq2 results to a signed per-feature weight vector:

where is the DESeq2 test statistic and is the nominal p-value.

### Gene-level gene set enrichment analysis (GSEA)

We performed gene-level pathway enrichment using FGSEA[49] on curated MSigDB gene-set collections (v2024.1). For each dataset, we concatenated Hallmark, GO Biological Process, Reactome, and KEGG collections and did not de-duplicate gene sets across collections.

Genes were ranked using the DESeq2-derived signed weights defined in. This ranking preserves the global sign convention, such that positive weights (and thus NES > 0) correspond to genes higher in the case condition.

Enrichment was computed with FGSEA. We report enrichment scores (ES), normalized enrichment scores (NES), and Benjamini-Hochberg adjusted q-values (FDR) computed over all tested gene sets within each dataset. Gene sets with FDR < 0.05 were considered significant for downstream Theme consolidation, and complete GSEA outputs (including leading-edge genes) are provided in Supplementary Table S7.

### Enhancer-program enrichment analysis (EPEA)

We performed enhancer-program enrichment analysis (EPEA) by applying FGSEA to enhancer-level (GHID) differential signals, treating enhancers as ranked features and curated enhancer programs as set definitions. Enhancers were ranked using the DESeq2-derived signed weights defined, preserving the global sign convention such that NES > 0 indicates enrichment among enhancers higher in the case condition.

For each EPEA run, we loaded the distributed base enhancer libraries and derived the analysis-time libraries by applying the single-pass specificity/size filter (Fig. 1a, parameters in Supplementary Table S1). For total RNA-seq datasets, we evaluated both cell-type and TFBS-defined libraries. Because canonical enhancer RNAs are predominantly non-polyadenylated and unstable [15], poly(A)-selected libraries attenuate enhancer-level signal; for mRNA/poly(A) inputs, enhancer-set collections were therefore evaluated separately as a sensitivity-oriented rule and combined at the reporting stage.

We computed Benjamini-Hochberg adjusted q-values (FDR) over the concatenated set list within each dataset. Unless stated otherwise, enhancer sets with q < 0.05 (total RNA-seq) or q < 0.10 (mRNA/poly(A) RNA-seq) were considered significant EPEA hits for downstream helper definition and robustness analyses. Complete EPEA outputs, including leading-edge GHIDs for each enriched set, are provided in Supplementary Table S8.

### Helper selection, robustness, and specificity

For total RNA-seq datasets, we defined candidate helpers as significant EPEA hits (q-value < 0.05) from the concatenated cell-type and TFBS-defined EPEA results. Candidate hits were pooled across libraries, ranked by −log10(q-value), and a Kneedle-style elbow was used to select the top *k* helpers (reported in Supplementary Table S2). For mRNA/poly(A) inputs, helpers were defined by a sensitivity-oriented rule as all significant EPEA hits under the mRNA configuration (q < 0.10) without elbow selection and without additional specificity filtering. Subsequently, using GeneHancer’s gene-enhancer associations, we identified genes associated with these enhancers to perform preliminary functional annotation of the helper using Hallmark pathways ORA.

To assess robustness of helper readouts, we performed leave-one-replicate-out (LORO) analyses within each dataset. For each fold, we removed one replicate, refit differential expression, recomputed signed weights, reran EPEA with the same analysis-time libraries and thresholds, and repeated helper selection under the same rules above. For the iPSC-Neuron dataset, which uses a donor-blocked design (∼ cell_donor + condition) across five donors, four of which contribute samples to both conditions, each of the 15 LORO folds removed a single sample rather than an entire donor; the design matrix remained rank-full across all folds because the remaining donors always spanned both conditions. Robustness was summarized as helper recurrence and sign consistency across folds (Supplementary Table S3). In the main text and Fig. 4b–c, we report robustness and specificity results for the subset of helpers linked to final Supported or Partial claims (20 helpers across four datasets); full per-helper results for all selected helpers are provided in Supplementary Tables S3–S4.

To test whether helper enrichments reflected specific set identity beyond set size, we generated size-matched null enhancer sets and compared observed helper enrichment to a shuffle-based null. For each dataset, we first computed observed EPEA statistics for the selected helpers (NES_obs). We then defined a sampling universe U as the union of GHIDs appearing in the analysis-time library concepts and generated S shuffled concept lists by, for each helper, sampling without replacement a random set of GHIDs from U with the same cardinality as the observed helper (no matching beyond set size).

Using the fixed observed weight vector, we reran EPEA on each shuffled concept list to obtain a null distribution of enrichment scores for each helper. A directional empirical p-value was computed in the observed direction as:

where is the number of non-missing null draws and is the indicator function.

For a standardized effect size, we computed, where and are the mean and standard deviation of helpers with were treated as ineligible for z-based summaries. We harmonized z to the observed direction as and combined evidence across eligible helpers using an equal-weight Stouffer method:

,

where is the number of eligible helpers and is the standard normal CDF. Minimum null-depth requirements and other thresholds are reported in Supplementary Table S1; outputs are summarized in Supplementary Table S4 and Fig. S4.

For a small number of helpers, the size-matched shuffle produced too few valid null enrichment scores for a reliable specificity test, because most randomly sampled size-matched sets were non-significant and returned no enrichment statistic (fewer than 50 finite nulls; Supplementary Table S4). This affected the BT20 mesenchymal stem cell program (3 valid nulls), *SUPT5H* in PANC1, and four helpers in iPSC-Neuron. For these helpers the observed EPEA enrichment is reported, but the shuffle-based specificity score is not interpretable; their helper status rests on EPEA significance and selection rather than the shuffle.

We additionally assessed null behavior under case/control label exchange for datasets where sample size permitted adequate permutation depth. For BT20-*BCL2L14-ETV6*-Fusion (P = 1,000 random permutations) and iPSC-Neuron (P = 1,000 random samples from C(15, 9) = 5,005 possible labelings), we refit DESeq2, recomputed weights, and reran EPEA for each permutation to obtain a null distribution of NES for each helper. For iPSC - Neuron, permutations were drawn globally (unrestricted) rather than stratified within donors; because the DESeq2 model (∼ cell_donor + condition) was refit for each permutation, donor effects were re-estimated under every label assignment, and all 1,000 permutations produced rank-full design matrices. This unrestricted scheme is conservative relative to within-donor stratification, as it admits label assignments that partially confound donor and condition effects, broadening the null distribution. Empirical p-values were computed from the full permutation distribution in the observed direction (Fig. 4d). Label permutation was not performed for PANC1-Rapa-1h (n = 6 samples; insufficient combinatorial depth) or MDA231-*SNAI1*-KO (n = 9 samples; exhaustive enumeration yielded only 84 unique permutations, reaching the combinatorial ceiling).

### Theme consolidation by data-driven clustering

We condensed significant GSEA gene sets into process-level Themes using a deterministic clustering pipeline. Significant gene sets (FDR < 0.05) were split by direction. Within each direction, pairwise Jaccard similarity was computed on leading-edge genes, and gene sets were clustered by average-linkage hierarchical clustering followed by adaptive tree cutting (dynamicTreeCut [50], with parameters auto-selected from input size; Supplementary Methods). Unassigned gene sets were retained as singleton orphans.

Each cluster was labelled by its medoid (the member with the highest mean Jaccard to other members; ties broken by q-value). When any cluster member matched one of approximately 30 curated canonical-process terms, this label was promoted over the medoid-derived label to preserve named minority signals. For each Theme, we reported the mean NES as the Theme effect, and a union of up to 12 leading-edge genes. A post-clustering merger then collapsed near-redundant Themes whose full leading-edge gene unions (computed from the complete GSEA output, not the 12-gene summary) exceeded a Jaccard threshold of 0.5, with Hallmark-anchored Themes retained as representatives.

### Evidence classification and verdict assignment

The upstream GSEA and EPEA outputs, together with the clustered Theme summaries, constitute a self-contained analytical result that can be interpreted directly by domain experts. The following evidence classification step is an optional interpretation aid that organizes these precomputed statistics using broad biological knowledge; it does not add new data or recompute any statistic. We classified each Theme by the strength of its enhancer-program support using a structured evidence classifier operating on precomputed Theme summaries and Helper metadata (Fig. 1c). No differential expression or enrichment statistics are recomputed at this stage.

A direction-matching gate first restricts consideration to Helpers sharing the Theme’s enrichment direction. For each direction-matched pair, the classifier assigns one of three verdict tiers: SUPPORTED, when the Helper’s regulatory identity is the defining feature of the Theme’s biological process; PARTIAL, when the Theme describes a characteristic but non-defining activity of the Helper’s program; or GENE_LEVEL_ONLY, when no specific mechanistic connection exists. Each Theme may link up to 3 Helpers, and each Helper may appear in at most 5 Themes.

Classification was implemented via Anthropic Claude Opus (temperature = 0) under a fixed prompt and deterministic output validation; outputs failing validation were auto-corrected for capacity violations or replaced by a conservative all-GENE_LEVEL_ONLY fallback (Supplementary Methods). For each SUPPORTED or PARTIAL verdict, we computed an evidence weight:

where is the EPEA q-value of each linked Helper. PARTIAL verdicts receive a 0.5× discount.

Each dataset was run three times and reconciled by majority verdict with Helper intersection across agreeing runs; unstable verdicts or empty intersections were conservatively assigned GENE_LEVEL_ONLY. All locked parameters, the verbatim classifier prompt, and per-dataset audit artefacts are deposited in the project repository.

### Enhancer over-representation analysis for GWAS traits (eORA)

We performed an enhancer-based over-representation analysis (eORA) to connect GWAS loci to cell-type enhancer programs (Fig. 5a). GWAS associations were obtained from the NHGRI-EBI GWAS Catalog and processed using the pruning/QC strategy described in the CATlas [34,51]: within each (trait, PMID), genome-wide significant associations were ranked by p-value and retained iteratively such that no two retained SNPs were within a small genomic window on the same chromosome. We then restricted to well-powered studies by requiring a minimum number of retained lead loci and a minimum reported sample size (cases for binary traits, or total participants for quantitative traits). For each trait, lead loci were pooled across PMIDs.

Lead SNP identifiers were normalized to rsIDs and converted to genomic coordinates on GRCh38 using Ensembl variation resources, retaining autosomes. SNPs were mapped to GeneHancer enhancers by direct interval overlap against a GeneHancer GRCh38 BED reference; SNPs overlapping multiple enhancers contributed to all overlapping GHIDs. For each trait, the trait enhancer set was defined as the union of mapped enhancers, restricted to the GHID universe of the tested cell-type enhancer sets.

Cell-type programs were defined by the curated cell-type enhancer-set library (excluding cancer cell lines) without additional ubiquity filtering for this analysis. For each trait enhancer set and each cell-type program, we computed enrichment by a right-tailed hypergeometric test using as background the union of GHIDs across all tested cell-type programs. P-values were adjusted within each trait across all tested cell types using Benjamini-Hochberg to yield trait-wise FDR values (reported in Supplementary Table S9). For the focused trait scans in Fig. 5c, we additionally report a Bonferroni display threshold based on the number of tested cell types.

As a mapping/calibration diagnostic, we computed a size-matched null for each trait by repeatedly sampling random GHID sets of the same size from the background universe and recomputing hypergeometric p-values for each cell type; cell-type-specific null cutoffs derived from these permutations were recorded as calibration fields and used for sanity checks (Fig. S5), but were not used as the primary significance criterion.

## Availability of data and materials

The RNA-seq dataset for BT20 cells expressing *BCL2L14-ETV6* fusion variants is available from the Gene Expression Omnibus under accession GSE120919. The three additional validation datasets analyzed in this study are publicly available: MDA-MB-231 *SNAI1*-knockout RNA-seq under GEO accession GSE210870, PANC-1 rapamycin ribosome-profiling and RNA-seq under GEO accession GSE225683, and the iPSC-to-neuron differentiation time course under the Sequence Read Archive accession PRJNA596331.

The EnSEMBLE software, including the R package and the Python evidence-classifier agent, together with the locked configuration, gene-set libraries, agent prompts, and per-dataset outputs used in this study, is openly available at https://github.com/wangxlab/EnSEMBLE and archived at Zenodo (DOI: 10.5281/zenodo.20495674; release v2.2.0). The complete enhancer overlap enrichment (eORA) results are provided as Supplementary Data.

GeneHancer v5.24, used for enhancer-to-gene mapping, is proprietary and is distributed under license by GeneCards/LifeMap Sciences. It cannot be redistributed and is therefore not included in the software repository or the Zenodo archive; it is available to qualifying users directly from GeneCards (https://www.genecards.org/) under license. The enhancer-set libraries provided as Supplementary Data list only GeneHancer element identifiers (GHIDs), shared with permission from LifeMap Sciences, and contain no other GeneHancer data fields (no coordinates, target genes, or scores).

## Competing interests

X.-S.W. is the founder of CA-Genome Rx Inc. CA-Genome Rx Inc. was not involved in the design, execution, analysis, or interpretation of this study, and no products, patents, or intellectual property related to CA-Genome Rx Inc. are discussed or evaluated in this work. The remaining authors declare no competing interests.

## Funding Information

This work was supported by CDMRP Breast Cancer Research Program (BCRP) under Award Nos. HT9425-24-1-0150, HT9425-24-1-0850, and W81XWH-22-1-1036. These grants provided salary support only; the research described here was not a direct objective of the funded BCRP projects. Opinions, interpretations, conclusions, and recommendations are those of the authors.

## Authors’ contributions

L.Z. and A.G. contributed equally to this work. L.Z.: Conceptualization, Methodology, Software, Formal analysis, Investigation, Data curation, Visualization, Writing – original draft. A.G.: Methodology, Software, Visualization, Writing – review & editing. Y.W.: Data curation, Resources, Writing – review & editing. R.S.: Data curation, Resources, Writing – review & editing. B.L.: Data curation, Resources, Writing – review & editing. G.H.: Visualization, Writing – review & editing. X.-S.W.: Conceptualization, Methodology, Supervision, Software, Formal analysis, Investigation, Funding acquisition, Writing – review & editing. All authors read and approved the final manuscript.

## Acknowledgements

This research was supported in part by the University of Pittsburgh Center for Research Computing, RRID:SCR_022735, through the resources provided. Specifically, this work used the HTC cluster, which is supported by NIH award number S10OD028483. This work also used the Bridges-2 system at the Pittsburgh Supercomputing Center (PSC), supported by NSF award number ACI-1928147, through allocation MCB180037.

## Additional Files

**Additional file 1: Supplementary Methods.** Detailed descriptions of clustering parameter selection, deterministic validation rules, auto-correction procedure, three-run consensus policy, and iPSC-Neuron dataset processing caveats.

**Additional file 2: Supplementary Figures S1-S5.** Figure S1: Composition and size distribution of enhancer-set libraries before filtering. Figure S2: Two-step filtering reshapes enhancer-set libraries. Figure S3: Differential enhancer activity across perturbations. Figure S4: Per-helper specificity under set-identity shuffles. Figure S5: eORA input coverage, mapping sanity check, and null calibration.

**Additional file 3: Supplementary Table S1.** Parameter ledger. Complete list of pipeline parameters across all analysis stages (RNA-seq quantification and differential expression, enhancer-set library construction, GSEA/EPEA, helper selection, theme clustering, robustness testing, the LLM agent, and eORA), with values and applicable dataset types.

**Additional file 4: Supplementary Table S2.** Helper selection. All elbow-selected helpers across the four datasets, with EPEA NES, q-value, class, direction, and flags for claim-linkage, set-permutation eligibility, and label-permutation eligibility.

**Additional file 5: Supplementary Table S3.** Leave-one-replicate-out (LORO) stability. Per-helper summary (mean, standard deviation, and range of NES across folds; sign-stability) and per-fold NES values for all selected helpers.

**Additional file 6: Supplementary Table S4.** Set-identity shuffle specificity. Per-helper observed NES, number of valid size-matched nulls (B), directional z-score (Z_dir), empirical p-value, and eligibility, for all selected helpers across the four datasets.

**Additional file 7: Supplementary Table S5.** Claim and verdict table. All 132 themes with their verdict tier (Supported / Partial / Gene-level only), theme weight, linked helpers, consensus rule, and classifier rationale.

**Additional file 8: Supplementary Table S6.** Theme membership. All 132 themes with their constituent gene sets, direction, mean NES, and representative leading-edge genes.

**Additional file 9: Supplementary Table S7.** Full GSEA output. Complete gene-set enrichment results across the four datasets.

**Additional file 10: Supplementary Table S8.** Full EPEA output. Complete enhancer-set enrichment results across the four datasets.

**Additional file 11: Supplementary Table S9.** eORA results. FDR-significant trait-by-cell-type enhancer enrichment results (q_BH < 0.05). Full results for all tested combinations are provided as Supplementary Data.

**Supplementary Data 1**: Complete eORA results (eORA_full_results.csv; all tested trait-by-cell-type combinations) and the per-dataset agent inputs and outputs (structured input, verdicts, verdict tables, narrative summaries, and reports) for the four validation datasets.

**Supplementary Data 2**: Cell-type enhancer-set library (normal_CellType2Enhancer.gmt; 411 sets, GeneHancer element identifiers shared with permission from LifeMap Sciences).

**Supplementary Data 3**: TFBS-defined enhancer-set library (enhancer_sets_tfbs_GeneHancer_v5.24.gmt; 537 sets, GeneHancer element identifiers shared with permission from LifeMap Sciences).

## References

1. Subramanian A, Tamayo P, Mootha VK, Mukherjee S, Ebert BL, Gillette MA, et al. Gene set enrichment analysis: A knowledge-based approach for interpreting genome-wide expression profiles. Proceedings of the National Academy of Sciences. Proceedings of the National Academy of Sciences; 2005;102:15545–50. 10.1073/pnas.0506580102

2. Reimand J, Isserlin R, Voisin V, Kucera M, Tannus-Lopes C, Rostamianfar A, et al. Pathway enrichment analysis and visualization of omics data using g:Profiler, GSEA, Cytoscape and EnrichmentMap. Nat Protoc. Nature Publishing Group; 2019;14:482–517. 10.1038/s41596-018-0103-9

3. Kuleshov MV, Jones MR, Rouillard AD, Fernandez NF, Duan Q, Wang Z, et al. Enrichr: a comprehensive gene set enrichment analysis web server 2016 update. Nucleic Acids Res. 2016;44:W90–7. 10.1093/nar/gkw377

4. Huang DW, Sherman BT, Lempicki RA. Systematic and integrative analysis of large gene lists using DAVID bioinformatics resources. Nat Protoc. Nature Publishing Group; 2009;4:44–57. 10.1038/nprot.2008.211

5. Wu T, Hu E, Xu S, Chen M, Guo P, Dai Z, et al. clusterProfiler 4.0: A universal enrichment tool for interpreting omics data. The Innovation. 2021;2:100141. 10.1016/j.xinn.2021.100141

6. Lee S, Deng L, Wang Y, Wang K, Sartor MA, Wang X-S. IndepthPathway: an integrated tool for in-depth pathway enrichment analysis based on single-cell sequencing data. Bioinformatics. 2023;39:btad325. 10.1093/bioinformatics/btad325

7. Khatri P, Sirota M, Butte AJ. Ten Years of Pathway Analysis: Current Approaches and Outstanding Challenges. PLOS Computational Biology. Public Library of Science; 2012;8:e1002375. 10.1371/journal.pcbi.1002375

8. Ziemann M, Schroeter B, Bora A. Two subtle problems with overrepresentation analysis. Bioinformatics Advances. 2024;4:vbae159. 10.1093/bioadv/vbae159

9. Begley CG, Ellis LM. Raise standards for preclinical cancer research. Nature. Nature Publishing Group; 2012;483:531–3. 10.1038/483531a

10. Freedman LP, Cockburn IM, Simcoe TS. The Economics of Reproducibility in Preclinical Research. PLOS Biology. Public Library of Science; 2015;13:e1002165. 10.1371/journal.pbio.1002165

11. Lee S, Hu Y, Loo SK, Tan Y, Bhargava R, Lewis MT, et al. Landscape analysis of adjacent gene rearrangements reveals BCL2L14–*ETV6* gene fusions in more aggressive triple-negative breast cancer. Proceedings of the National Academy of Sciences. Proceedings of the National Academy of Sciences; 2020;117:9912–21. 10.1073/pnas.1921333117

12. Sabino AU, Guerreiro D de M, Kim A-R, Ramos AF, Reinitz J. Characterizing the regulatory logic of transcriptional control at the DNA sequence level by ensembles of thermodynamic models. Bioinformatics. 2025;41:btaf534. 10.1093/bioinformatics/btaf534

13. Arnold PR, Wells AD, Li XC. Diversity and Emerging Roles of Enhancer RNA in Regulation of Gene Expression and Cell Fate. Front Cell Dev Biol [Internet]. Frontiers; 2020 [cited 2026 Jan 26];7. 10.3389/fcell.2019.00377

14. Kaikkonen MU, Spann NJ, Heinz S, Romanoski CE, Allison KA, Stender JD, et al. Remodeling of the Enhancer Landscape during Macrophage Activation Is Coupled to Enhancer Transcription. Molecular Cell. Elsevier; 2013;51:310–25. 10.1016/j.molcel.2013.07.010

15. Li W, Notani D, Rosenfeld MG. Enhancers as non-coding RNA transcription units: recent insights and future perspectives. Nat Rev Genet. Nature Publishing Group; 2016;17:207–23. 10.1038/nrg.2016.4

16. Li W, Notani D, Ma Q, Tanasa B, Nunez E, Chen AY, et al. Functional roles of enhancer RNAs for oestrogen-dependent transcriptional activation. Nature. Nature Publishing Group; 2013;498:516–20. 10.1038/nature12210

17. Schaukowitch K, Joo J-Y, Liu X, Watts JK, Martinez C, Kim T-K. Enhancer RNA Facilitates NELF Release from Immediate Early Genes. Molecular Cell. Elsevier; 2014;56:29–42. 10.1016/j.molcel.2014.08.023

18. Pratapa A, Jalihal AP, Law JN, Bharadwaj A, Murali TM. Benchmarking algorithms for gene regulatory network inference from single-cell transcriptomic data. Nat Methods. Nature Publishing Group; 2020;17:147–54. 10.1038/s41592-019-0690-6

19. Aibar S, González-Blas CB, Moerman T, Huynh-Thu VA, Imrichova H, Hulselmans G, et al. SCENIC: single-cell regulatory network inference and clustering. Nat Methods. Nature Publishing Group; 2017;14:1083–6. 10.1038/nmeth.4463

20. Nguyen H, Tran D, Tran B, Pehlivan B, Nguyen T. A comprehensive survey of regulatory network inference methods using single cell RNA sequencing data. Brief Bioinform. 2021;22:bbaa190. 10.1093/bib/bbaa190

21. Upadhya SR, Ryan CJ. Experimental reproducibility limits the correlation between mRNA and protein abundances in tumor proteomic profiles. Cell Reports Methods [Internet]. Elsevier; 2022 [cited 2026 Jan 26];2. 10.1016/j.crmeth.2022.100288

22. McLean CY, Bristor D, Hiller M, Clarke SL, Schaar BT, Lowe CB, et al. GREAT improves functional interpretation of cis-regulatory regions. Nat Biotechnol. Nature Publishing Group; 2010;28:495–501. 10.1038/nbt.1630

23. Qin Q, Fan J, Zheng R, Wan C, Mei S, Wu Q, et al. Lisa: inferring transcriptional regulators through integrative modeling of public chromatin accessibility and ChIP-seq data. Genome Biol. 2020;21:32. 10.1186/s13059-020-1934-6

24. Sheffield NC, Bock C. LOLA: enrichment analysis for genomic region sets and regulatory elements in R and Bioconductor. Bioinformatics. 2016;32:587–9. 10.1093/bioinformatics/btv612

25. Layer RM, Pedersen BS, DiSera T, Marth GT, Gertz J, Quinlan AR. GIGGLE: a search engine for large-scale integrated genome analysis. Nat Methods. Nature Publishing Group; 2018;15:123–6. 10.1038/nmeth.4556

26. Welch RP, Lee C, Imbriano PM, Patil S, Weymouth TE, Smith RA, et al. ChIP-Enrich: gene set enrichment testing for ChIP-seq data. Nucleic Acids Res. 2014;42:e105. 10.1093/nar/gku463

27. Lee CT, Cavalcante RG, Lee C, Qin T, Patil S, Wang S, et al. Poly-Enrich: count-based methods for gene set enrichment testing with genomic regions. NAR Genom Bioinform. 2020;2:lqaa006. 10.1093/nargab/lqaa006

28. Schep AN, Wu B, Buenrostro JD, Greenleaf WJ. chromVAR: inferring transcription-factor-associated accessibility from single-cell epigenomic data. Nat Methods. Nature Publishing Group; 2017;14:975–8. 10.1038/nmeth.4401

29. Lee C, Wang K, Qin T, Sartor MA. Testing Proximity of Genomic Regions to Transcription Start Sites and Enhancers Complements Gene Set Enrichment Testing. Front Genet [Internet]. Frontiers; 2020 [cited 2026 Jul 6];11. 10.3389/fgene.2020.00199

30. Maurano MT, Humbert R, Rynes E, Thurman RE, Haugen E, Wang H, et al. Systematic Localization of Common Disease-Associated Variation in Regulatory DNA. Science. American Association for the Advancement of Science; 2012;337:1190–5. 10.1126/science.1222794

31. Nasser J, Bergman DT, Fulco CP, Guckelberger P, Doughty BR, Patwardhan TA, et al. Genome-wide enhancer maps link risk variants to disease genes. Nature. Nature Publishing Group; 2021;593:238–43. 10.1038/s41586-021-03446-x

32. Rao SSP, Huntley MH, Durand NC, Stamenova EK, Bochkov ID, Robinson JT, et al. A 3D Map of the Human Genome at Kilobase Resolution Reveals Principles of Chromatin Looping. Cell. Elsevier; 2014;159:1665–80. 10.1016/j.cell.2014.11.021

33. The GTEx Consortium. The GTEx Consortium atlas of genetic regulatory effects across human tissues. Science. American Association for the Advancement of Science; 2020;369:1318–30. 10.1126/science.aaz1776

34. Cerezo M, Sollis E, Ji Y, Lewis E, Abid A, Bircan KO, et al. The NHGRI-EBI GWAS Catalog: standards for reusability, sustainability and diversity. Nucleic Acids Res. 2025;53:D998–1005. 10.1093/nar/gkae1070

35. Tsirigoti C, Ali MM, Maturi V, Heldin C-H, Moustakas A. Loss of *SNAI1* induces cellular plasticity in invasive triple-negative breast cancer cells. Cell Death Dis. Nature Publishing Group; 2022;13:832. 10.1038/s41419-022-05280-z

36. Jägle S, Busch H, Freihen V, Beyes S, Schrempp M, Boerries M, et al. SNAIL1-mediated downregulation of FOXA proteins facilitates the inactivation of transcriptional enhancer elements at key epithelial genes in colorectal cancer cells. PLOS Genetics. Public Library of Science; 2017;13:e1007109. 10.1371/journal.pgen.1007109

37. Wu Y, Deng J, Rychahou PG, Qiu S, Evers BM, Zhou BP. Stabilization of Snail by NF-κB Is Required for Inflammation-Induced Cell Migration and Invasion. Cancer Cell. Elsevier; 2009;15:416–28. 10.1016/j.ccr.2009.03.016

38. Saxton RA, Sabatini DM. mTOR Signaling in Growth, Metabolism, and Disease. Cell. Elsevier; 2017;168:960–76. 10.1016/j.cell.2017.02.004

39. Rahl PB, Lin CY, Seila AC, Flynn RA, McCuine S, Burge CB, et al. c-Myc Regulates Transcriptional Pause Release. Cell. Elsevier; 2010;141:432–45. 10.1016/j.cell.2010.03.030

40. Baluapuri A, Hofstetter J, Stankovic ND, Endres T, Bhandare P, Vos SM, et al. MYC Recruits SPT5 to RNA Polymerase II to Promote Processive Transcription Elongation. Molecular Cell. Elsevier; 2019;74:674–687.e11. 10.1016/j.molcel.2019.02.031

41. Liberzon A, Birger C, Thorvaldsdóttir H, Ghandi M, Mesirov JP, Tamayo P. The Molecular Signatures Database Hallmark Gene Set Collection. cels. Elsevier; 2015;1:417–25. 10.1016/j.cels.2015.12.004

42. Nguyen TU, Hector H, Pederson EN, Lin J, Ouyang Z, Wendel H-G, et al. Rapamycin-Induced Feedback Activation of eIF4E-EIF4A Dependent mRNA Translation in Pancreatic Cancer. Cancers [Internet]. 2023 [cited 2026 Jan 27];15. 10.3390/cancers15051444

43. Burke EE, Chenoweth JG, Shin JH, Collado-Torres L, Kim S-K, Micali N, et al. Dissecting transcriptomic signatures of neuronal differentiation and maturation using iPSCs. Nat Commun. Nature Publishing Group; 2020;11:462. 10.1038/s41467-019-14266-z

44. Pauklin S, Vallier L. The Cell-Cycle State of Stem Cells Determines Cell Fate Propensity. Cell. Elsevier; 2013;155:135–47. 10.1016/j.cell.2013.08.031

45. Keenan AB, Torre D, Lachmann A, Leong AK, Wojciechowicz ML, Utti V, et al. ChEA3: transcription factor enrichment analysis by orthogonal omics integration. Nucleic Acids Res. 2019;47:W212–24. 10.1093/nar/gkz446

46. Imrichová H, Hulselmans G, Kalender Atak Z, Potier D, Aerts S. i-cisTarget 2015 update: generalized cis-regulatory enrichment analysis in human, mouse and fly. Nucleic Acids Res. 2015;43:W57–64. 10.1093/nar/gkv395

47. Hu M, Alkhairy S, Lee I, Pillich RT, Fong D, Smith K, et al. Evaluation of large language models for discovery of gene set function. Nat Methods. Nature Publishing Group; 2025;22:82–91. 10.1038/s41592-024-02525-x

48. Wang Z, Jin Q, Wei C-H, Tian S, Lai P-T, Zhu Q, et al. GeneAgent: self-verification language agent for gene-set analysis using domain databases. Nat Methods. Nature Publishing Group; 2025;22:1677–85. 10.1038/s41592-025-02748-6

49. Korotkevich G, Sukhov V, Budin N, Shpak B, Artyomov MN, Sergushichev A. Fast gene set enrichment analysis [Internet]. bioRxiv; 2021 [cited 2026 Jan 27]. p. 060012. 10.1101/060012

50. Langfelder P, Zhang B, Horvath S. Defining clusters from a hierarchical cluster tree: the Dynamic Tree Cut package for R. Bioinformatics. 2008;24:719–20. 10.1093/bioinformatics/btm563

51. Zhang K, Hocker JD, Miller M, Hou X, Chiou J, Poirion OB, et al. A single-cell atlas of chromatin accessibility in the human genome. Cell. 2021;184:5985–6001.e19. 10.1016/j.cell.2021.10.024

